# eDNA metabarcoding vs metagenomics: an assessment of dietary competition in two estuarine pipefishes

**DOI:** 10.1101/2021.01.05.425398

**Authors:** Conny P. Serite, Arsalan Emami-Khoyi, Ofentse K. Ntshudisane, Nicola C. James, Bettine Jansen van Vuuren, Taryn Bodill, Paul D. Cowley, Alan K. Whitfield, Peter R. Teske

**Affiliations:** Centre for Ecological Genomics and Wildlife Research, Department of Zoology, University of Johannesburg, Auckland Park 2006, South Africa; National Research Foundation – South African Institute for Aquatic Biodiversity, Private Bag 1015, Makhanda 6140, South Africa

**Keywords:** DNA barcoding, environmental DNA (eDNA), fecal DNA, diet analysis, endangered species

## Abstract

Understanding the dietary preferences of endangered species can be useful in implementing conservation strategies, including habitat restoration, translocation, and captive breeding. Environmental DNA (eDNA) from feces provides a non-invasive method for analyzing animal diets. Currently, metabarcoding, a PCR-based method, is the method of choice for analyzing such data. However, this method has limitations, specifically PCR bias, which can result in the overestimation of the importance of certain taxa and the failure to detect other taxa because they do not amplify. The present study compared metabarcoding with metagenomics, a marker-free method, to assess the diversity of prey items in the feces of a critically endangered South African estuarine pipefish, *Syngnathus watermeyeri*, and its widely distributed congener (*S. temminckii*) to investigate the potential dietary competition. Our results showed a distinct difference between the diets of *S. watermeyeri* and *S. temminckii*, with the former mainly consuming calanoid copepods and the latter preferring caridean shrimp. Metagenomics produced more species identifications than metabarcoding; however, most of the species identified are not present in South Africa. The identifications made by both methods mostly belonged to the same families, but some prey items were identified only by either method. Both methods would benefit from a complete reference database of South African estuarine macroinvertebrates.

## Introduction

Estuaries are amongst the most threatened aquatic habitats in the world, many of which have become functionally degraded due to anthropogenic pressures (Edgar et al., 2000; Turpie et al., 2002; Kaselowski and Adams, 2013; Kajee et al., 2018). These include water abstraction for agricultural and industrial activities, pollution, and urban development (Orth et al., 2006). As such, some storage reservoirs in water-scarce countries such as South Africa now have the potential to retain more than 50% of the freshwater that the estuaries would receive under normal conditions (Wooldridge and Callahan, 2000). Several endemic estuarine species in South Africa are threatened, including the Endangered Knysna seahorse, *Hippocampus capensis* (Lockyear et al., 2006; Mkare et al., 2017), the Critically Endangered limpet *Siphonaria compressa* (Allanson and Herbert, 2005) and the Critically Endangered estuarine pipefish, *Syngnathus watermeyeri* (Whitfield, 1995). All three species are associated with submerged macrophyte beds mainly dominated by the eelgrass *Zostera capensis*, which is itself listed as vulnerable by the IUCN because it is sensitive to the current level of anthropogenic pressure and experiences widespread degradation as a result of increased coastal development (Payne et al., 1998; Adams, 2016). With declines in their natural habitat, ecosystem restoration, translocation, and captive breeding need to be considered as a means to conserve the remaining populations of endangered species (Strum, 2005; Gumm et al., 2011; Landa et al., 2017). Therefore, a thorough knowledge of what these species consume in the wild is required to provide ecosystem managers with the information necessary to manage these populations better. Reconstruction of diet in wild populations is critical in ecology because it reveals important details about a species’ feeding habits (Pompanon et al., 2012), how a species uses its surroundings, and if there is resource competition with other members of the same community (Klare et al., 2011; Mumma et al., 2016). Animal diets have traditionally been determined by morphological examination of the gastrointestinal and fecal contents or by direct observation of their feeding habits (Pompanon et al., 2012; Sousa et al., 2019; Harper et al., 2020). However, the hard-part remains of some prey items can be difficult to identify using morphological analysis since they are usually damaged beyond recognition (Hawlitschek et al., 2018; Harper et al., 2020). In addition, studying the diet of endangered species can be challenging particularly when the species of interest is rare and elusive, making field studies especially difficult (Ang et al., 2010).

A recently developed alternative method of assessing animal diets is eDNA metabarcoding (Kartzinel et al., 2015; Srivathsan et al., 2015; Boukhdoud et al., 2021). This PCR-based approach amplifies short DNA fragments of specified genetic markers that can then be identified using known reference sequences (Shehzad et al., 2012). However, metabarcoding has shortcomings, including PCR bias (Ferravante et al., 2021); this may occur due to irregular primer binding, thus resulting in some species amplifying worse than others or not at all (Alberdi et al., 2018; Mata et al., 2019). Metagenomics is an alternative approach that involves the direct random sequencing of the entire genomic DNA rather than a small number of genetic markers (Bohmann et al., 2014; Bovo et al., 2018; Piñol, 2021). One of its drawbacks is that the bulk of the DNA that has been sequenced cannot be reliably assigned taxonomic rank due to a lack of comprehensive reference sequences since, for most species, only small portions of the entire genome have so far been sequenced (Bovo et al., 2018; Piñol, 2021).

Here, we compared the use of metagenomics and metabarcoding in identifying the prey species found in the feces of the estuarine pipefish and compared the prey items identified with corresponding data from its sister species, the longsnout pipefish *S. temminckii*, which is more abundant and widely distributed. Since the two pipefish share the same habitats and both capture small prey items by expanding their buccal cavity and suctioning prey through their tubular snouts (van Wassenbergh et al., 2008), it was hypothesized that dietary competition might exist between the two (Whitfield et al., 2017).

## Materials and methods

### Study sites and sample collection

Collection of *Syngnathus watermeyeri* and *S. temminckii* was approved by the Department of Agriculture, Forestry and Fisheries of South Africa (permit: RES2018/107), and ethical clearance was granted by the SAIAB Animal Ethics Committee (reference number: 25/4/1/5_2018-07) and the University of Johannesburg Faculty of Science Ethics Committee (reference number: 2021-10-05/Serite_Teske). Samples were collected from the only two South African estuaries where the estuarine pipefish still occurs, the Bushmans and Kariega (Claassens et al., 2022; Weiss et al., 2022). Pipefishes were collected at three sites; two in the Bushmans Estuary (site 1: 33°40’43.9 “S 26°39’12.2”E; site 2: 33°40’21.6”S 26°38’46.1”E; data from these sites was subsequently pooled) and the third site in the Kariega Estuary (33°39’10.4”S 26°39’04.6”E).

Sampling was conducted between March 30^th^ and April 6^th^, 2019. In each location, all the pipefishes that could be collected within a period of 2 hours using a 5 mm stretch mesh seine net were placed into 5ℓ plastic tanks containing estuarine water for 3 h, after which they were released back into the estuaries. A total of 13 *S. watermeyeri* and 29 *S. temminckii* specimens were collected and kept in tanks in small groups of 2-3 individuals. The tanks were kept in the shade and aerated using portable air pumps, and the water was replaced every 30 min. Fecal pellets were dropped by the pipefishes in all these tanks throughout the 3 h period, and were immediately collected using a sterile Pasteur pipette for each species, and subsequently blotted dry by placing them on paper towels before preserving them into 2 ml screw-cap microcentrifuge tubes containing RNAlater stabilization and storage reagent (QIAGEN GmbH, Hilden, Germany). The tubes were kept frozen on ice for up to two days and then stored at −70°C upon returning to the laboratory.

### Laboratory analysis

Prior to DNA extraction, the fecal pellets were thawed at room temperature and then transferred to new 1.5ml microcentrifuge tubes, which were placed on a heat block for 2 h at 37°C. DNA extraction was done in triplicate for each sample using three different extraction protocols, and the best samples were selected for sequencing: the CTAB procedure (Doyle, 1991), as well as two extraction kit methods, NucleoSpin and Qiagen, following manufacturers’ instructions. For metabarcoding, the mitochondrial cytochrome oxidase c subunit I (COI) gene was amplified using forward primer mlCOIintF and reverse primer jgHCO2198 (Leray et al., 2013) as described in Ntuli et al. (2020). The PCR products were purified using the AMPure XP system (Beckman Coulter), and a NEBNext Ultra DNA Library Prep Kit (New England BioLabs, United States) was used for the preparation of genomic libraries. The resulting libraries were screened for size distribution using a 2100 Bioanalyzer (Agilent) and quantified using real-time PCR. The libraries were then sequenced on an Illumina HiSeq 4000 platform (Illumina Inc., San Diego, California, United States) at Novogene (Hong Kong), using 2X250 bp paired-end chemistry according to the manufacturer’s instructions.

For metagenomics, 0.4 μg of genomic DNA was used for library preparations. The libraries were generated using a NEBNext DNA Library Prep Kit (New England BioLabs, United States), and indices were then added to each sample. The genomic DNA was randomly sheared into fragments of 350 bp. The fragments were end-polished, A-tailed, and ligated using the NEBNext adapter for Illumina sequencing, and the fragments were PCR enriched by P5 and indexed P7 oligos. The PCR products were purified using the AMPure XP system (Beckman Coulter), and the resulting libraries were screened for size distribution using a 2100 Bioanalyzer (Agilent) and quantified using real-time PCR. Genomic libraries were sequenced on an Illumina Novaseq6000 platform (Illumina Inc., San Diego, California, United States) at Novogene (Hong Kong), using 2X150 bp paired-end chemistry according to the manufacturer’s instructions.

### Sequence assembly and analysis

For the metabarcoding, quality control was carried out using FastQC (http://www.bioinformatics.bbsrc.ac.uk/projects/fastqc/) and sequencing adapters, all sequences with length less than 150 bp, and low-quality sequences, which were defined as those sequences with a quality Phred Score of less than 25 in a five bp sliding window, were removed using Trimmomatic v0.36 (Bolger et al., 2014) Then, Cutadapt v4.1 (Martin, 2011) was used to trim both forward and reverse primer sequences. When only the forward or reverse read of a particular sequence passed the quality filtering step, the expected error rate for “unpaired forward” and “unpaired reverse” was estimated in VSEARCH v2.17.0 (Rognes et al., 2016), and since the forward sequences consistently showed a lower error rate compared to reverse sequences, only a subset of full length (250 bp) forward sequences were selected for downstream analysis, together with sequences produced by merging forward and reverse reads.

Metabarcoding sequences were assembled using VSEARCH v2.17.0 pipeline. Briefly, all pair-end sequences were merged based on their overlaps and merged sequences, and full-length only forward sequences were separately dereplicated into unique sequences. Chimeric amplicons were removed using a denovo method implemented in the same package, and all non-chimeric sequences with a minimum of 98% similarity were clustered into distinct groups, also known as operational taxonomic units (or OTUs). The consensus sequence for each cluster and the number of sequences that formed each cluster were extracted for the taxonomic rank assignment step.

The metagenomic sequences from each location were separately assembled into longer contigs using MEGAHIT v1.1.1 (Li et al., 2015) by selecting the “meta-large” preset, which is most appropriate for complex metagenomic assemblies (https://github.com/voutcn/megahit). When possible, assembled sequences were dereplicated using VMATCH (Kurtz, 2003), and the quality of the assemblies was assessed with QUAST v4.6.3 (Gurevich et al., 2013).

### Taxonomic rank assignment

To assign a taxonomic rank to consensus sequences, metabarcoding sequences were blast-searched (Altschul et al., 1990) against a local non-redundant COI database, using a minimum similarity score of 98% and a minimum query coverage of 150 bp. Then, assembled contigs from metagenomics were blast-searched against the complete NCBI nucleotide database ftp://ftp.ncbi.nlm.nih.gov/blast/db/.

A consensus taxonomic rank was assigned to each sequence based on the Last Common Ancestor (LCA) of the five best matches, as explained in BASTA (Kahlke and Ralph, 2019). Non-target taxa (i.e., contaminants, pipefish DNA, and taxa that are too large or small to constitute prey, including humans, bacteria, and algae) were bioinformatically removed. The OTUs were then subset based on classes that contributed more than 1% to the overall read counts. The OTU counts for each species were then agglomerated into the taxonomic rank of class and were subsequently normalized relative to the total OTU counts estimated for each species using the phyloseq R package v1.38.0 (McMurdie and Holmes, 2013). The final results were visualized in ggplot2 v3.3.5 (Wickham, 2016). A list of the potential prey items identified by metabarcoding and metagenomics was compiled. The prior information about the presence or absence of the identified species in South African estuaries was checked using the Global Biodiversity Information Facility (GBIF) database.

## Results

The metabarcoding sequencing run generated 6 442 764 sequences from the Bushmans Estuary and 8 706 470 sequences from the Kariega Estuary for *S. watermeyeri*. For *S. temminckii*, 7 803 100 sequences and 8 540 218 sequences were recovered for the Bushmans Estuary and the Kariega Estuary, respectively. Post-quality filtering, the number of paired-end sequences kept per sample ranged from 1 197 706 to 1 972 358 (Supplementary Table 1), and 23 005 to 77 265 for the forward sequences (Supplementary Table 2). 30 510 OTUs were generated for the metabarcoding sequences, and 712 were assigned a taxonomic rank, 649 were used for downstream analysis.

The metagenomic assemblies generated 611 473 (N50 = 707) and 183 631 (N50 = 2 952) singleton contigs for *S. watermeyeri* from the Bushmans and Kariega estuaries, respectively. For *S. temminckii*, 467 665 (N50 = 650) singleton contigs were generated for the Kariega Estuary samples and 514 739 (N50 = 720) for the Bushmans Estuary samples (Supplementary Table 3). Of 1819 of the overall sequences assigned a taxonomic rank, 408 were used for downstream analysis.

Metabarcoding results for each species showed a clear preference of each pipefish for a specific prey class (Figure 1). The estuarine pipefish preferred species in class Hexanauplia, with 90% of the OTUs belonging to this class, whereas 90% of the OTUs in *S. temminckii* belong to class Malacostraca (Supplementary Table 4). All of the Hexanauplia recovered sequences in the feces of *S. watermeyeri* were identified as a single species, *Pseudodiaptomus hessei;* similarly, close to 100% of the Malacostraca reads in *S. temminckii* were identified as *Palaemon peringueyi* (<0.002% matched to four other species, Table 1). Gastropods were found to be a noticeable dietary component in *S. watermeyeri* only (2%, Figure 1, Supplementary Table 5), with ~90% of the sequences identified as *Assiminea capensis*), ~10% as *Hydrobia knysnaensis* and < 1% as *Afrolittorina africana*. Only one sequence in the diet of *S. temminckii* was identified as a gastropod, *Bursatella leachi*. Among a total of 11 potential prey item species reconstructed from metabarcoding of feces, nine species occur in South Africa.

**FIGURE 1:**
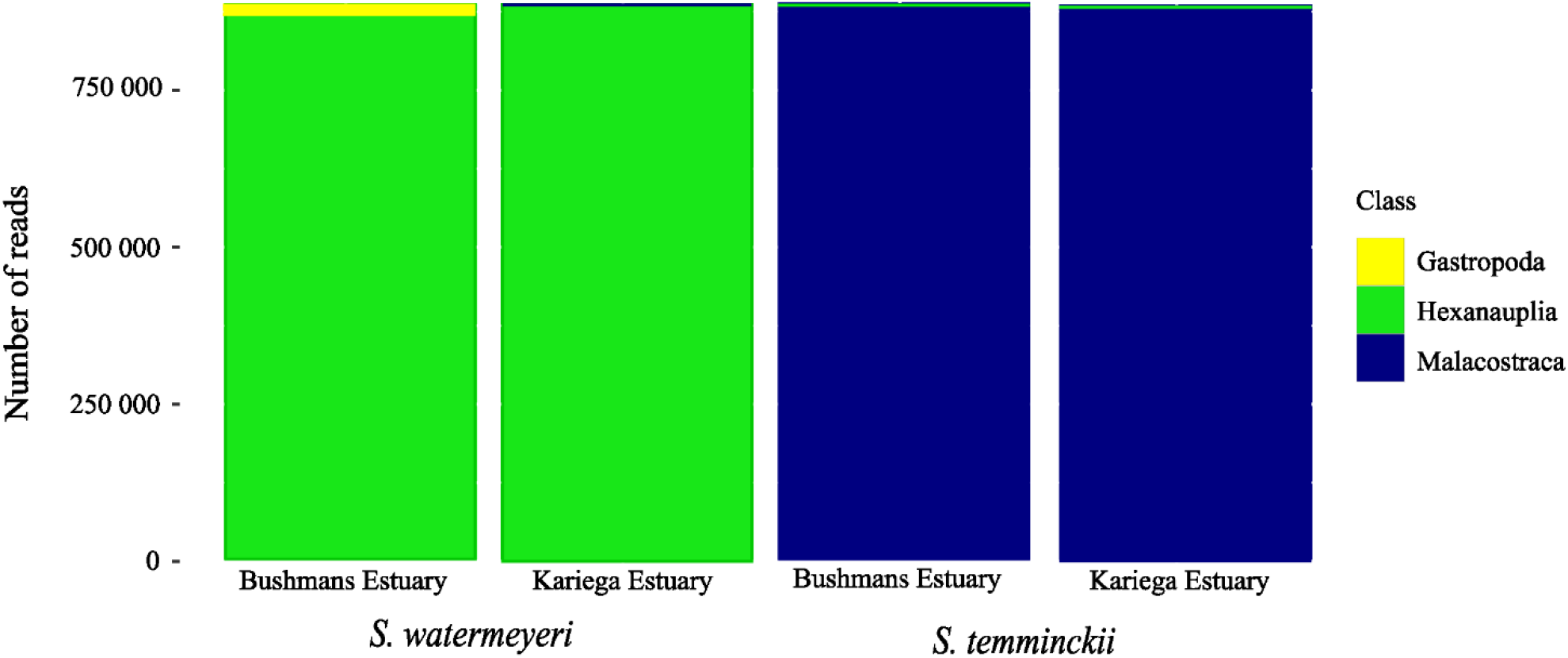
Comparison of the read abundance of three prey classes (Gastropoda, Hexanauplia, and Malacostraca) found in the metabarcoding data generated using feces of the two pipefish species. The figure shows a subset of prey classes that contributed more than 1% of the total read abundance per pipefish sample.

**TABLE 1.** Prey species associated with classes found in the feces of both *S. watermeyeri* (W) and *S. temminckii* (T) based on metabarcoding and metagenomics. Black squares indicate the presence of a taxon, and superscripts represent ^K^Kariega Estuary only; BBushmans Estuary only; *species present in South Africa; ^?^genus present in South Africa.

| Class | Species | Metabarcoding |  | Metagenomics |  |
| --- | --- | --- | --- | --- | --- |
|  |  | W | T | W | T |
| Arachnida | *Medioppia subpectinata* |  |  | ■ B |  |
|  | *Oppiella nova* * |  |  | ■ B |  |
| Bivalvia | *Perna perna* * | ■ K |  |  |  |
| Gastropoda | *Afrolittorina africana* * | ■ B |  |  |  |
|  | *Assiminea capensis* * | ■ B |  | ■ B |  |
|  | *Bursatella leachii* * |  | ■ B |  |  |
|  | *Hydrobia knysnaensis* * | ■ B |  |  |  |
|  | *Oncomelania hupensis* |  |  | ■ B |  |
|  | *Patella pellucida* ? |  |  |  | ■ B |
| Hexanauplia | *Halicyclops* sp. |  |  | ■ |  |
|  | *Paracalanus parvus* |  | ■ B |  |  |
|  | *Phyllodiaptomus praedictus* |  |  | ■ K |  |
|  | *Pseudodiaptomus euryhalinus* ? |  |  | ■ B |  |
|  | *Pseudodiaptomus hessei* * | ■ | ■ | ■ K |  |
|  | *Pseudodiaptomus nihonkaiensis* ? |  |  | ■ K |  |
| Insecta | *Liposcelis brunnea* ? |  | ■ B |  |  |
|  | *Platycorypha nigrivirga* |  | ■ K |  |  |
|  | *Psorophora longipalpus* | ■ K |  |  |  |
|  | *Tuberculatus annulatus* * |  | ■ K |  |  |
| Malacostraca | *Balsscallichirus gilchristi* * | ■ K |  |  |  |
|  | *Crangon franciscorum* ? |  |  |  | ■ |
|  | *Exosphaeroma hylecoetes* * |  | ■ B |  |  |
|  | *Homarus americanus* |  |  |  | ■ K |
|  | *Macrobrachium nipponense* ? |  |  |  | ■ |
|  | *Macrobrachium rosenbergii* ? |  |  |  | ■ |
|  | *Palaemon carinicauda* ? |  |  |  | ■ |
|  | *Palaemon elegans* ? |  |  |  | ■ |
|  | *Palaemon modestus* ? |  |  |  | ■ K |
|  | *Palaemon pacificus* ? |  |  |  | ■ K |
|  | *Palaemon paludosus* ? |  |  |  | ■ |
|  | *Palaemon peringueyi* * | ■ K | ■ |  | ■ |
|  | *Palaemon pugio* ? |  |  |  | ■ B |
|  | *Palaemon serenus* ? |  |  |  | ■ |
|  | *Palaemon serratus* ? |  |  |  | ■ |
|  | *Palaemon tonkinensis* ? |  |  |  | ■ K |
|  | *Palaemon varians* ? |  |  |  | ■ B |
|  | *Penaeus chinensis* ? |  |  |  | ■ B |
|  | *Penaeus indicus* * |  | ■ K |  |  |
|  | *Portumnus* sp. | ■ K | ■ |  |  |
| Ostracoda | *Cyprideis torosa* * |  |  |  | ■ B |
|  | *Darwinula stevensoni* * |  |  | ■ B |  |
| Polychaeta | *Phylo* sp. * | ■ K | ■ K |  |  |
|  | *Prionospio* sp. * | ■ |  |  |  |
| Thecostraca | *Sacculina carcini* |  |  | ■ B |  |

Metagenomics recovered more species than metabarcoding (Table 1). This was particularly true within the Malacostraca, where 16 species were identified by metagenomics and only five by metabarcoding, with *Palaemon peringueyi* being the only species in common. However, most of the potential prey species identified by the metagenomic method do not occur in South Africa; of the 24 species identified, only three are found in the study area: *P. peringueyi, P. hessei*, and *A. capensis*. Additionally, both methods reported three additional prey classes not identified by the other. Metabarcoding identified species from classes Bivalvia, Insecta, and Polychaeta, collectively contributing 3% of the total OTUs for *S. watermeyeri* and 2% for *S. temminckii* (Polychaeta and Insecta only). Metagenomics identified species from Arachnida, Ostracoda, and Thecostraca, most of these identifications were for *S. watermeyeri*, with one species, *Cyprideis torosa*, being identified in the diet of *S. temminckii*.

## Discussion

This study used metagenomic, and metabarcoding analysis of eDNA collected non-invasively from fecal samples to reconstruct and compare the dietary preferences of the Critically Endangered estuarine pipefish, *Syngnathus watermeyeri*, and its more abundant congener *S. temminckii*. The research aimed to assess the capabilities of both methods in dietary assessment and provide practical information that contributes towards improving the conservation management of S. watermeyeri, particularly in captive breeding and releases of captive-bred progeny into estuaries within the historical range.

The results showed clear differences in dietary preferences between the two pipefish species. Although some species were found in both species, each pipefish had a clear preference for one prey class. Due to the high conservation status of the estuarine pipefish, fecal samples could only be collected from a few individuals, and for financial reasons, fecal material from each species and estuary was pooled. Therefore, this study provides a snapshot of diet composition in each population rather than information on individual preferences. However, the fact that all individuals from the same site were captured on the same day indicates that both species had access to the same prey species, thus rejecting the idea that the differences in diet composition could be due to small sample sizes or spatial and temporal separation of the captured individuals.

While it is possible that species from the largely terrestrial classes Arachnida and Insecta (few of which have larvae that can survive in estuarine water), as well as classes that were identified using only one of the two methods (e.g., Bivalvia, Ostracoda, Polychaeta, etc.), are opportunistically consumed or represent secondary predation by one of the prey items, only two invertebrate classes (Malacostraca and Hexanauplia) were consistently found in the diet of the two pipefish species, and in each case, these were limited to very few species. The estuarine pipefish has a preference for copepods (class Hexanauplia). The calanoid copepod *Pseudodiaptomus hessei* dominated the metabarcoding reads and was also found using metagenomics. In contrast, although *S. temminckii* also consume this copepod, this pipefish shows a preference for a caridean shrimp (class Malacostraca), *Palaemon peringueyi*. In addition to crustaceans, both metabarcoding and metagenomics confirmed the importance of the gastropod *Assimenia capensis* in the diet of the estuarine pipefish. As the adults of Palaemonidae and *Assimenia* are too large to fit through the mouths of pipefishes, it is likely that larvae are preyed on.

As is the case in all PCR-based approaches, our results concerning the importance of these main prey items based on metabarcoding may have been impacted by amplification bias (Tedersoo et al., 2015; Krehenwinkel et al., 2017). However, given that the primers used here have a higher success rate in amplifying the DNA of metazoan species present in fecal samples than any other primers currently in use (Leray et al., 2013), and that the species that made up the bulk of the metabarcoding reads were also recovered using metagenomics, suggests that amplification bias cannot account for the clear difference in the number of OTUs amplified for potential prey species whose DNA was present in the feces of both pipefish species.

Metagenomics is predicted to provide superior taxonomic resolution compared to metabarcoding due to its capacity to use multiple markers across the genome and to assemble longer contigs for more accurate species identification (Srivathsan et al., 2016; Chua et al., 2021). However, this depends on how comprehensive the reference database is regarding the species represented and the markers used to amplify them (Chua et al., 2021). As seen in this study, most metagenomic identifications were for species not present in the study area; this was particularly evident in the class Malacostraca. In contrast to the metabarcoding reads, most of which originated from *Palaemon peringueyi*, several additional caridean shrimps of the family Palaemonidae (which includes the genera *Palaemon* and *Macrobrachium*) were identified using metagenomics, and this is likely an artifact of gene regions being represented in this dataset that has not yet been sequenced for *P. peringueyi* and consequently the taxonomic rank of a closely related congener with a complete genome were reported. Hence, both sequencing methods used in this study will display bias due to incomplete databases when there is a lack of diverse species representation in reference databases (Chua et al., 2021).

The differences in prey selection between the two pipefish species may depend on clear differences in snout length and gape size, as well as variations in hunting tactics. The estuarine pipefish’s shorter snout size may restrict the size of prey species on which it can feed, which would explain its preference for small copepods and gastropod veliger larvae, as well as ostracods (metagenomic data only). It is also possible that *S. watermeyeri* may detect small zooplankton more effectively than *S. temminckii* because of the position of its eyes, which are located closer to the tip of the snout. In addition to stronger suction and greater gape size due to a larger and longer snout, *S. temminckii* has been observed to hunt its prey actively, whereas *S. watermeyeri* is more passive and waits to ambush prey animals to swim within reach (Sven-Erick Weiss, pers. obs.). Together, these factors likely allow *S. temminckii* to stalk prey that might otherwise escape (such as the larvae of *P. peringueyi*). However, it does not solely rely on these prey items.

The finding that the Critically Endangered *S. watermeyeri* relies to a large extent on relatively small zooplankters (particularly copepods) supports the hypothesis that significant reductions in zooplankton abundance in response to reduced freshwater influx (Grange et al., 2000) can result in estuarine pipefish population declines (Whitfield et al., 2017). Excessive freshwater abstraction has transformed both estuaries inhabited by *S. watermeyeri* from systems with well-developed salinity gradients to homogeneously marine-dominated systems. The Bushmans River, for example, has approximately 30 impoundments in its upper reaches that have significantly reduced freshwater inflow (Bornman and Klages, 2004). While this would have resulted in a decrease in phytoplankton biomass (and, by extension, zooplankton biomass) (Hilmer and Bate, 1990), the resulting increase in water clarity would also have facilitated the formation of the current extensive submerged macrophyte beds (Bornman and Klages, 2004). Because of this contradiction (reduced food availability but increased habitat availability), it cannot be ruled out that the two estuaries in their current marine-dominated state have a higher carrying capacity for *S. watermeyeri* than they would have had under natural conditions, at least during periods of moderate rainfall.

The large number of OTUs from the calanoid copepod *P. hessei* in the feces of *S. watermeyeri* was clearly linked to low freshwater inflow into this estuary during the study period. The zooplankton community of the Bushmans Estuary has been poorly studied, but during low or zero river flow periods in the Kariega Estuary, *P. hessei* dominates the zooplankton community (up to 76% by number), this can change following river flooding when another calanoid, *Acartia longipatella*, becomes more abundant (Froneman and Vorwerk, 2013). Possible dietary switches by *S. watermeyeri* according to changes in copepod species composition in the zooplankton community have yet to be determined.

This study suggests that resource competition between the two pipefish species is likely negligible as they mostly consume a different diet. Thus, it is not an important factor that could explain why *S. watermeyeri* is such a rare species. Although the estuarine pipefish prefers copepods, both sequencing methods indicate that its diet can be supplemented in captivity with gastropod veliger larvae to resemble how they would eat in their natural environment. Other taxa may also be opportunistically consumed, including insects or arachnids that may have fallen into the water and bivalve larvae, ostracods, and polychaetes. Overall, eDNA sequencing methods have offered a way of identifying soft-bodied prey that might have been difficult or even impossible to detect through morphological analysis of fecal samples, such as the copepods and polychaetes identified here, while the identification of veliger shells in the feces would have been insufficient to identify the species from which they originated. However, the results also document how the lack of reference sequences can greatly impact the use of metabarcoding and metagenomics as diet analysis methods. Our study highlights the critical role of a local genome-wide reference database for more accurate reconstruction of dietary items from environmental samples and the need for a more comprehensive reference database of South African estuarine macroinvertebrates.

## Author Contributions

PC, AW, and PT conceived the research. PC and PT generated funding and obtained permits. ON, AE-K, PC, NJ, TB, and PT participated in sampling. CS and AE-K analyzed the data. CS created figures and tables. AE-K, BJvV, and PT supervised the students. All authors contributed to the article and approved the submitted version.

## Acknowledgments

The Centre for High-Performance Computing (CHPC) is thanked for granting us access to the Lengau Cluster, and the University of Johannesburg IT services are acknowledged for their support. CPS is grateful to the Department of Higher Education for awarding her the Nurturing Emerging Scholars Program (NESP) scholarship. Sven-Erick Weiss, Jody-Carynn Oliver, Laura Tensen, and Claudia Schnelle are thanked for their assistance during sampling and laboratory analysis.

## Data Availability Statement

The raw read datasets generated and analyzed in this study will be submitted to the NCBI Sequence Read Archive (SRA) repository

## SUPPLEMENTARY INFORMATION

**Table 1:**
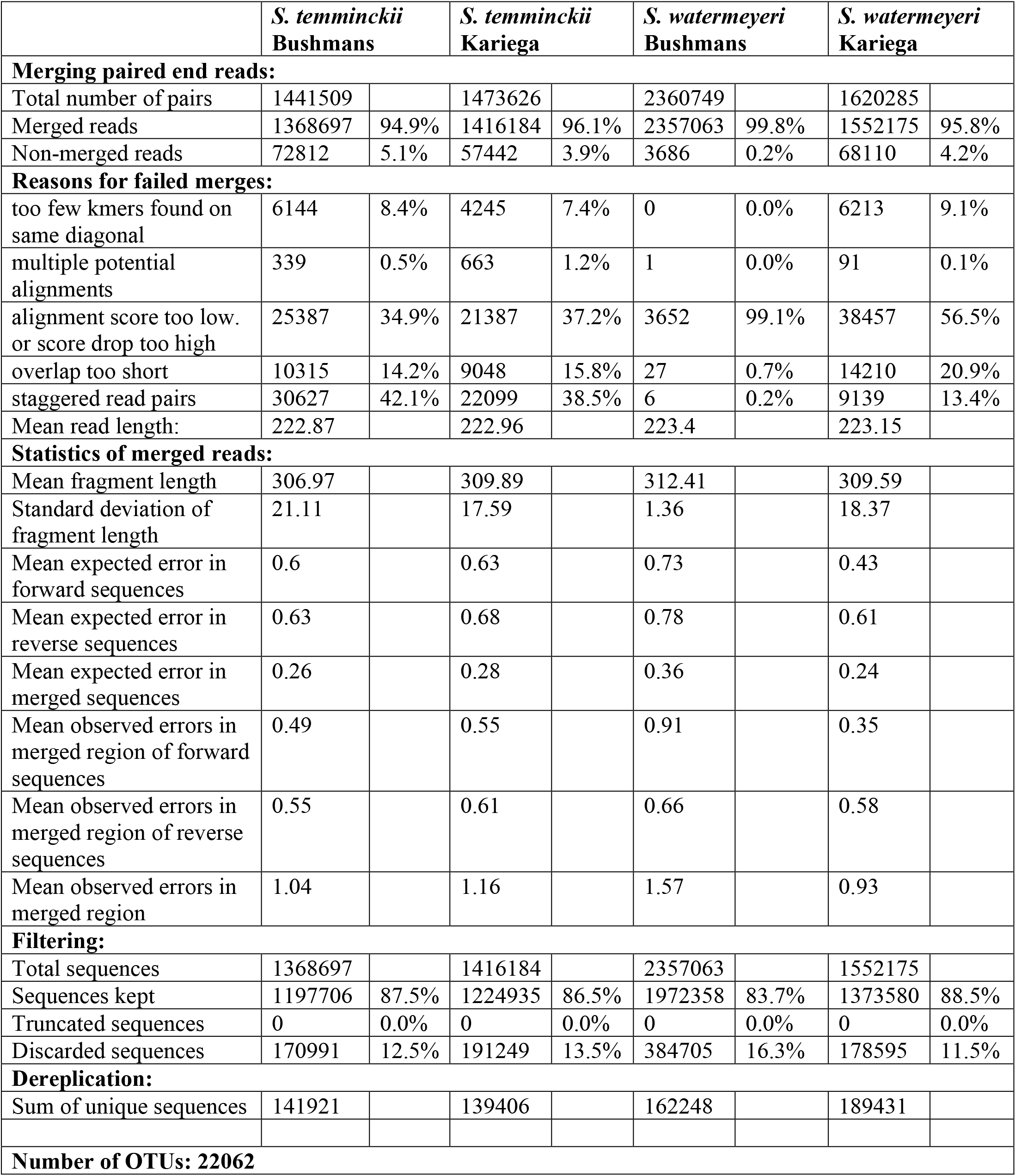
Metabarcoding paired end read filtering statistics

|  | *S. temminckii*<br>Bushmans |  | *S. temminckii*<br>Kariega |  | *S. watermeyeri*<br>Bushmans |  | *S. watermeyeri*<br>Kariega |  |
| --- | --- | --- | --- | --- | --- | --- | --- | --- |
| Merging paired end reads: |  |  |  |  |  |  |  |  |
| Total number of pairs | 1441509 |  | 1473626 |  | 2360749 |  | 1620285 |  |
| Merged reads | 1368697 | 94.9% | 1416184 | 96.1% | 2357063 | 99.8% | 1552175 | 95.8% |
| Non-merged reads | 72812 | 5.1% | 57442 | 3.9% | 3686 | 0.2% | 68110 | 4.2% |
| Reasons for failed merges: |  |  |  |  |  |  |  |  |
| too few kmers found on same diagonal | 6144 | 8.4% | 4245 | 7.4% | 0 | 0.0% | 6213 | 9.1% |
| multiple potential alignments | 339 | 0.5% | 663 | 1.2% | 1 | 0.0% | 91 | 0.1% |
| alignment score too low. or score drop too high | 25387 | 34.9% | 21387 | 37.2% | 3652 | 99.1% | 38457 | 56.5% |
| overlap too short | 10315 | 14.2% | 9048 | 15.8% | 27 | 0.7% | 14210 | 20.9% |
| staggered read pairs | 30627 | 42.1% | 22099 | 38.5% | 6 | 0.2% | 9139 | 13.4% |
| Mean read length: | 222.87 |  | 222.96 |  | 223.4 |  | 223.15 |  |
| Statistics of merged reads: |  |  |  |  |  |  |  |  |
| Mean fragment length | 306.97 |  | 309.89 |  | 312.41 |  | 309.59 |  |
| Standard deviation of fragment length | 21.11 |  | 17.59 |  | 1.36 |  | 18.37 |  |
| Mean expected error in forward sequences | 0.6 |  | 0.63 |  | 0.73 |  | 0.43 |  |
| Mean expected error in reverse sequences | 0.63 |  | 0.68 |  | 0.78 |  | 0.61 |  |
| Mean expected error in merged sequences | 0.26 |  | 0.28 |  | 0.36 |  | 0.24 |  |
| Mean observed errors in merged region of forward sequences | 0.49 |  | 0.55 |  | 0.91 |  | 0.35 |  |
| Mean observed errors in merged region of reverse sequences | 0.55 |  | 0.61 |  | 0.66 |  | 0.58 |  |
| Mean observed errors in merged region | 1.04 |  | 1.16 |  | 1.57 |  | 0.93 |  |
| Filtering: |  |  |  |  |  |  |  |  |
| Total sequences | 1368697 |  | 1416184 |  | 2357063 |  | 1552175 |  |
| Sequences kept | 1197706 | 87.5% | 1224935 | 86.5% | 1972358 | 83.7% | 1373580 | 88.5% |
| Truncated sequences | 0 | 0.0% | 0 | 0.0% | 0 | 0.0% | 0 | 0.0% |
| Discarded sequences | 170991 | 12.5% | 191249 | 13.5% | 384705 | 16.3% | 178595 | 11.5% |
| Dereplication: |  |  |  |  |  |  |  |  |
| Sum of unique sequences | 141921 |  | 139406 |  | 162248 |  | 189431 |  |
| Number of OTUs: 22062 |  |  |  |  |  |  |  |  |

**Table 2:** Metabarcoding single end read filtering statistic

|  | *S. temminckii*<br>Bushmans | *S. temminckii*<br>Kariega | *S. watermeyeri*<br>Bushmans | *S. watermeyeri*<br>Kariega |
| --- | --- | --- | --- | --- |
| **Filtering:** |  |  |  |  |
| Total sequences | 214 651 | 249 411 | 121 082 | 177 332 |
| Sequences kept | 77 265 | 75 706 | 23 005 | 51 899 |
| Truncated sequences | 0 | 0 | 0 | 0 |
| Discarded sequences | 137 386 | 173 705 | 98 077 | 125 433 |
| **Dereplication:** |  |  |  |  |
| Sum of unique sequences | 20 798 | 19 995 | 6 917 | 18 951 |
| **Number of OTUs: 8448** |  |  |  |  |

**Table 3:** Metagenomic assembly QUAST report

|  | *S. watermeyeri* |  | *S. temminckii* |  |
| --- | --- | --- | --- | --- |
| **Assembly** | **Kariega** | **Bushmans** | **Kariega** | **Bushmans** |
| **Total number of contigs** | 611 473 | 183 631 | 467 665 | 514 739 |
| **Number of contigs (<= 0 bp)** | 2 015 987 | 781 019 | 2 268 677 | 2 318 770 |
| **Number of contigs (<= 1000 bp)** | 68 572 | 95 837 | 30 429 | 6 1371 |
| **Number of contigs (<= 5000 bp)** | 366 | 12 277 | 56 | 2 411 |
| **Number of contigs (<= 10000 bp)** | 72 | 1 527 | 4 | 625 |
| **Number of contigs (<= 25000 bp)** | 8 | 51 | 0 | 87 |
| **Number of contigs (<= 50000 bp)** | 2 | 3 | 0 | 11 |
| **Total length (<= 0 bp)** | 926 358 012 | 530 247 357 | 906 243 861 | 991 982 362 |
| **Total length (<= 1000 bp)** | 94 397 322 | 280 060 512 | 39 702 280 | 115 255 374 |
| **Total length (<= 5000 bp)** | 3 279 286 | 91 856 688 | 369 189 | 22 892 484 |
| **Total length (<= 10000 bp)** | 1 333 696 | 20 726 945 | 54 386 | 10 869 062 |
| **Total length (<= 25000 bp)** | 431 170 | 1 702 565 | 0 | 3 135 536 |
| **Total length (<= 50000 bp)** | 227 331 | 156 729 | 0 | 642 530 |
| **Largest contig** | 117 579 | 55 180 | 16 608 | 76 115 |
| **Total length** | 449 225 316 | 338 168 213 | 317 506 061 | 407 025 308 |
| **GC (%)** | 35.94 | 43.84 | 37.21 | 39.59 |
| **N50** | 707 | 2 952 | 650 | 720 |
| **N75** | 584 | 1 414 | 561 | 586 |
| **L50** | 227 381 | 32 638 | 185 425 | 168 126 |
| **L75** | 403 126 | 73 631 | 317 452 | 326 300 |
| **Number of Ns per 100 kbp** | 0 | 0 | 0 | 0 |
| All statistics are based on contigs of size <= 500 bp. unless otherwise noted (e.g., "# contigs (<= 0 bp)" and "Total length (<= 0 bp)" include all contigs). |  |  |  |  |

**Table 4:** Total number of OTUs used for data interpretation and the number of OTUs in each sample per site

| Class | # OTUs | *S. watermeyeri* |  | *S. temminckii* |  |
| --- | --- | --- | --- | --- | --- |
|  |  | Bushmans | Kariega | Bushmans | Kariega |
| **Arachnida** | 2<br>(<1%) | 0<br>(0%) | 2<br>(<1%) | 0<br>(0%) | 0<br>(0%) |
| **Bivalvia** | 1<br>(<1%) | 0<br>(0%) | 1<br>(0,3%) | 0<br>(0%) | 0<br>(0%) |
| **Gastropoda** | 13<br>(2%) | 12<br>(7,7%) | 0<br>(0%) | 1<br>(<1%) | 0<br>(0%) |
| **Hexanauplia** | 313<br>(48,2%) | 139<br>(89,7%) | 274<br>(93,5%) | 17<br>(8%) | 17<br>(6,9%) |
| **Insecta** | 8<br>(1%) | 2<br>(1,3%) | 1<br>(<1%) | 1<br>(<1%) | 5<br>(2%) |
| **Malacostraca** | 309<br>(47,6%) | 0<br>(0%) | 12<br>(4,1%) | 193<br>(91%) | 224<br>(90,7%) |
| **Polychaeta** | 3<br>(<1%) | 2<br>(1,3%) | 3<br>(1%) | 0<br>(0%) | 1<br>(<1%) |
|  | **649** | **155** | **293** | **212** | **247** |

**Table 5:**
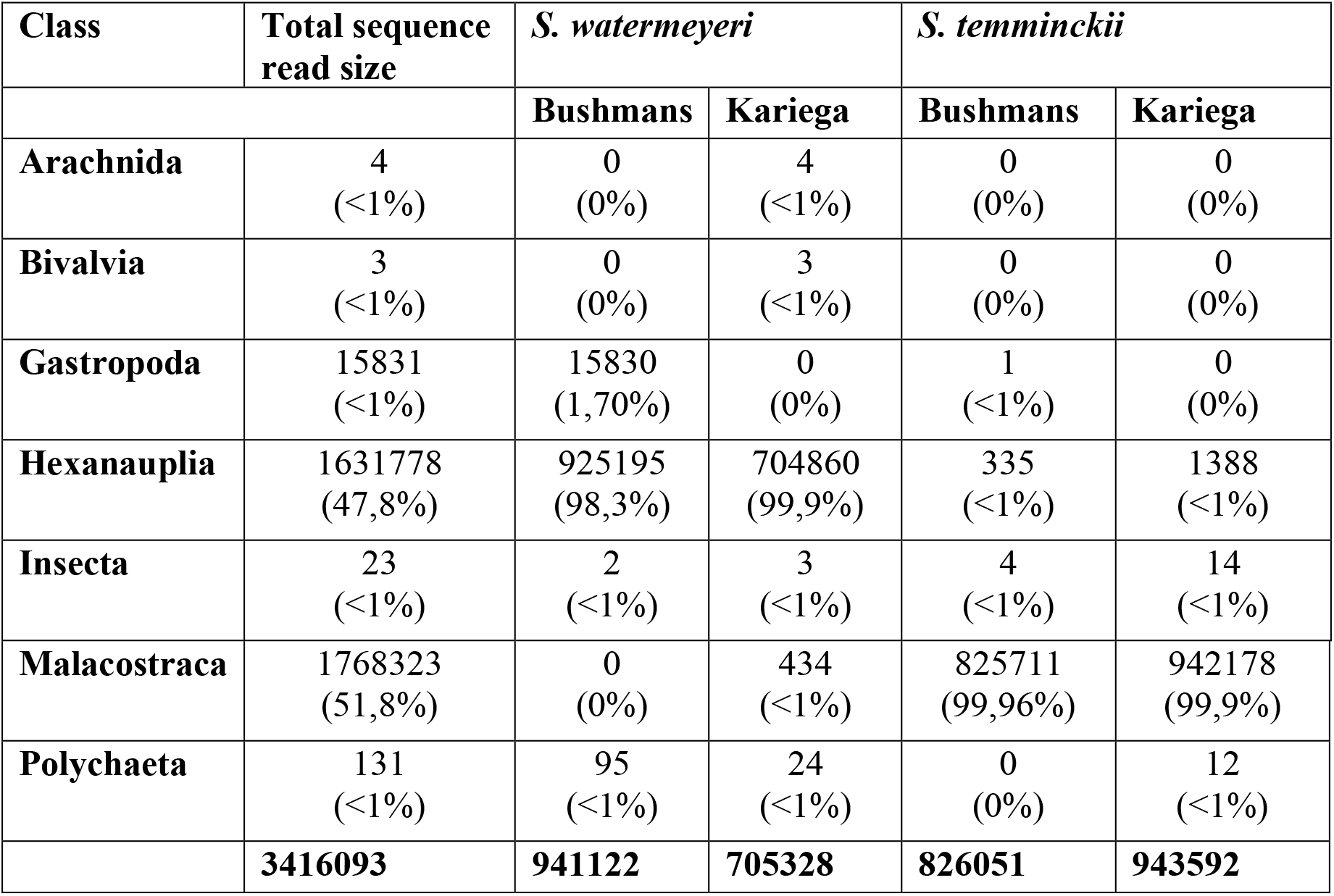
Number of sequences reads per class used for final analysis with the corresponding read sizes of each class

**Table 6:**
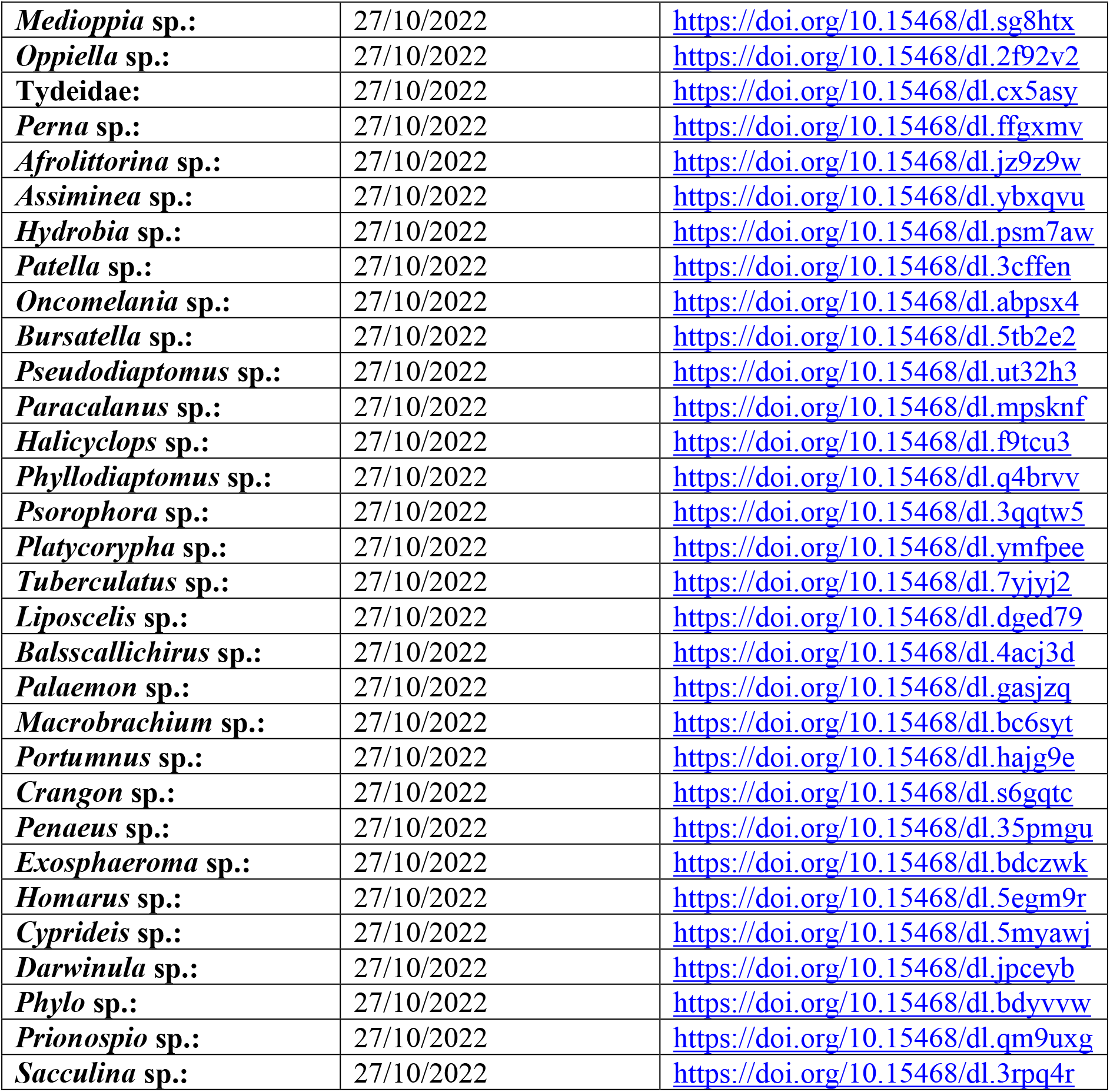
GBIF occurrence data download links

## References

Adams, J. B. (2016). Distribution and status of *Zostera capensis* in South African estuaries — A review. S Afr J Bot 107, 63–73. doi: 10.1016/j.sajb.2016.07.007.

Alberdi, A., Aizpurua, O., Gilbert, M. T. P., and Bohmann, K. (2018). Scrutinizing key steps for reliable metabarcoding of environmental samples. Methods Ecol Evol 9, 134–147. doi: 10.1111/2041-210X.12849.

Allanson, B. R., and Herbert, D. G. (2005). A newly discovered population of the critically endangered false limpet *Siphonaria compressa* Allanson, 1958 (Pulmonata: Siphonariidae), with observations on its reproductive biology. S Afr J Sci 101, 95–97. doi: 10.10520/EJC96338.

Altschul, S. F., Gish, W., Miller, W., Myers, E. W., and Lipman, D. J. (1990). Basic local alignment search tool. J Mol Biol 215, 403–410. doi: 10.1016/S0022-2836(05)80360-2.

Ang, A., Ismail, M. R. B., and Meier, R. (2010). Reproduction and infant pelage colouration of the banded leaf monkey (Mammalia: Primates: Cercopithecidae) in Singapore. Raffles Bull Zool 58, 411–415.

Bohmann, K., Evans, A., Gilbert, M. T. P., Carvalho, G. R., Creer, S., Knapp, M., et al. (2014). Environmental DNA for wildlife biology and biodiversity monitoring. Trends Ecol Evol 29, 358–367. doi: 10.1016/j.tree.2014.04.003.

Bolger, A. M., Lohse, M., and Usadel, B. (2014). Trimmomatic: a flexible trimmer for Illumina sequence data. Bioinformatics 30, 2114–2120. doi: 10.1093/bioinformatics/btu170.

Bornman, T., and Klages, N. (2004). Upgrade of Kenton-on-Sea/Bushmans River Mouth by Albany Coast Water Board.

Boukhdoud, L., Saliba, C., Parker, L. D., McInerney, N. R., Kahale, R., Saliba, I., et al. (2021). Using DNA metabarcoding to decipher the diet plant component of mammals from the Eastern Mediterranean region. Metabarcoding Metagenom 5, 219–231. doi: 10.3897/mbmg.5.70107.

Bovo, S., Ribani, A., Utzeri, V. J., Schiavo, G., Bertolini, F., and Fontanesi, L. (2018). Shotgun metagenomics of honey DNA: evaluation of a methodological approach to describe a multi-kingdom honey bee derived environmental DNA signature. PLoS One 13, e0205575. doi: 10.1371/journal.pone.0205575.

Chua, P. Y. S., Crampton-Platt, A., Lammers, Y., Alsos, I. G., Boessenkool, S., and Bohmann, K. (2021). Metagenomics: A viable tool for reconstructing herbivore diet. Mol Ecol Resour 21, 2249–2263. doi: 10.1111/1755-0998.13425.

Claassens, L., de Villiers, N. M., Seath, J., and Wasserman, J. (2022). Distribution and density of the Critically Endangered estuarine pipefish across its range – implications for conservation. Aquat Conserv 32, 28–41. doi: 10.1002/aqc.3742.

Doyle, J. (1991). “DNA protocols for plants,” in Molecular techniques in taxonomy (Springer Berlin Heidelberg), 283–293. doi: 10.1007/978-3-642-83962-7_18.

Edgar, G. J., Barrett, N. S., Graddon, D. J., and Last, P. R. (2000). The conservation significance of estuaries: A classification of Tasmanian estuaries using ecological, physical and demographic attributes as a case study. Biol Conserv 92, 383–397. doi: 10.1016/S0006-3207(99)00111-1.

Ferravante, C., Memoli, D., Palumbo, D., Ciaramella, P., di Loria, A., D’Agostino, Y., et al. (2021). HOME-BIO (shotgun metagenomic analysis of biological entities): a specific and comprehensive pipeline for metagenomic shotgun sequencing data analysis. BMC Bioinformatics 22, 106. doi: 10.1186/s12859-021-04004-y.

Froneman, P. W., and Vorwerk, P. D. (2013). Response of the plankton to a fresh water pulse in a fresh water deprived, permanently open South African estuary. J Water Resour Prot 05, 405–413. doi: 10.4236/jwrap.2013.54040.

Grange, N., Whitfield, A. K., de Villiers, C. J., and Allanson, B. R. (2000). The response of two South African east coast estuaries to altered river flow regimes. Aquat Conserv 10, 155–177. doi: 10.1002/1099-0755(200005/06)10:3<155::AID-AQC406>3.0.CO;2-Z.

Gumm, J. M., Snekser, J. L., Leese, J. M., Little, K. P., Leiser, J. K., Imhoff, V. E., et al. (2011). Management of interactions between endangered species using habitat restoration. Biol Conserv 144, 2171–2176. doi: 10.1016/J.biocon.2011.05.006.

Gurevich, A., Saveliev, V., Vyahhi, N., and Tesler, G. (2013). QUAST: quality assessment tool for genome assemblies. Bioinformatics 29, 1072–1075. doi: 10.1093/bioinformatics/btt086.

Harper, L. R., Watson, H., Donnelly, R., Hampshire, R., Sayer, C. D., Breithaupt, T., et al. (2020). Using DNA metabarcoding to investigate diet and niche partitioning in the native European otter (*Lutra lutra*) and invasive American mink (*Neovison vison*). Metabarcoding Metagenom 4, 113–133. doi: 10.3897/mbmg.4.56087.

Hawlitschek, O., Fernández-González, A., Balmori-de la Puente, A., and Castresana, J. (2018). A pipeline for metabarcoding and diet analysis from fecal samples developed for a small semi-aquatic mammal. PLoS One 13, 1–20. doi: 10.1371/journal.pone.0201763.

Hilmer, T., and Bate, G. C. (1990). Covariance analysis of chlorophyll distribution in the Sundays River Estuary, Eastern Cape. South Afri J Aquat Sci 16, 37–59. doi: 10.1080/10183469.1990.10557366.

Kahlke, T., and Ralph, P. J. (2019). BASTA–Taxonomic classification of sequences and sequence bins using last common ancestor estimations. Methods Ecol Evol 10, 100–103. doi: 10.1111/2041-210X.13095.

Kajee, M., Griffiths, C. L., and Lamberth, S. J. (2018). Long-term physico-chemical and faunal changes in a small, rural South African estuary. Afr Zool 53, 127–137. doi: 10.1080/15627020.2018.1552838.

Kartzinel, T. R., Chen, P. A., Coverdale, T. C., Erickson, D. L., Kress, W. J., Kuzmina, M. L., et al. (2015). DNA metabarcoding illuminates dietary niche partitioning by African large herbivores. Proc Natl Acad Sci U S A 112, 8019–8024. doi: 10.1073/pnas.1503283112.

Kaselowski, T., and Adams, J. B. (2013). Not so pristine-characterising the physico-chemical conditions of an undescribed temporarily open/closed estuary. Water SA 39, 627–636. doi: 10.4314/wsa.v39i5.6.

Klare, U., Kamler, J. F., and Macdonald, D. (2011). A comparison and critique of different scat-analysis methods for determining carnivore diet. Mamm Rev 41, 294–312. doi: 10.1111/j.1365-2907.2011.00183.x.

Krehenwinkel, H., Wolf, M., Lim, J. Y., Rominger, A. J., Simison, W. B., and Gillespie, R. G. (2017). Estimating and mitigating amplification bias in qualitative and quantitative arthropod metabarcoding. Sci Rep 7, 1–12. doi: 10.1038/s41598-017-17333-x.

Kurtz, S. (2003). The Vmatch large scale sequence analysis software. Computer Program, 4–12.

Landa, A., Flagstad, Ø., Areskoug, V., Linnell, J. D. C., Strand, O., Ulvund, K. R., et al. (2017). The endangered Arctic fox in Norway—the failure and success of captive breeding and reintroduction. Polar Res 36. doi: 10.1080/17518369.2017.1325139.

Leray, M., Yang, J. Y., Meyer, C. P., Mills, S. C., Agudelo, N., Ranwez, V., et al. (2013). A new versatile primer set targeting a short fragment of the mitochondrial COI region for metabarcoding metazoan diversity: application for characterizing coral reef fish gut contents. Front Zool 10, 34. doi: 10.1186/1742-9994-10-34.

Li, D., Liu, C.-M., Luo, R., Sadakane, K., and Lam, T.-W. (2015). MEGAHIT: an ultra-fast single-node solution for large and complex metagenomics assembly via succinct *de Bruijn* graph. Bioinformatics 31, 1674–1676. doi: 10.1093/bioinformatics/btv033.

Lockyear, J., Hecht, T., Kaiser, H., and Teske, P. (2006). The distribution and abundance of the endangered Knysna seahorse *Hippocampus capensis* (Pisces: Syngnathidae) in South African estuaries. Afr J Aquat Sci 31, 275–283.

Mata, V. A., Rebelo, H., Amorim, F., McCracken, G. F., Jarman, S., and Beja, P. (2019). How much is enough? Effects of technical and biological replication on metabarcoding dietary analysis. Mol Ecol 28, 165–175. doi: 10.1111/mec.14779.

McMurdie, P., and Holmes, S. (2013). Phyloseq: an R package for reproducible interactive analysis and graphics of microbiome census data. PLoS One 8, e61217.

Mkare, T. K., van Vuuren, B. J., and Teske, P. R. (2017). Conservation implications of significant population differentiation in an endangered estuarine seahorse. Biodivers Conserv 26, 1275–1293. doi: 10.1007/s10531-017-1300-5.

Mumma, M. A., Adams, J. R., Zieminski, C., Fuller, T. K., Mahoney, S. P., and Waits, L. P. (2016). A comparison of morphological and molecular diet analyses of predator scats. J Mammal 97, 112–120. doi: 10.1093/jmammal/gyv160.

Ntuli, N. N., Nicastro, K. R., Zardi, G. I., Assis, J., McQuaid, C. D., and Teske, P. R. (2020). Rejection of the genetic implications of the “Abundant Centre Hypothesis” in marine mussels. Sci Rep 10, 1–12. doi: 10.1038/s41598-020-57474-0.

Orth, R. J., Carruthers, T. J. B., Dennison, W. C., Duarte, C. M., Fourqurean, J. W., Heck, K. L., et al. (2006). A global crisis for seagrass ecosystems. Bioscience 56, 987–996. doi: 10.1641/0006-3568(2006)56[987:AGCFSE]2.0.CO;2.

Payne, M. F., Rippingale, R. J., and Longmore, R. B. (1998). Growth and survival of juvenile pipefish *Stigmatopora argus* fed live copepods with high and low HUFA content. Aquaculture 167, 237–245.

Piñol, J. (2021). Genotype by sequencing: An alternative new method to amplicon metabarcoding and shotgun metagenomics for the assessment of eukaryote biodiversity. Mol Ecol Resour 00, 1–4. doi: 10.1111/1755-0998.13320.

Pompanon, F., Deagle, B. E., Symondson, W. O. C., Brown, D. S., Jarman, S. N., and Taberlet, P. (2012). Who is eating what: Diet assessment using next generation sequencing. Mol Ecol 21, 1931–1950. doi: 10.1111/j.1365-294X.2011.05403.x.

Rognes, T., Flouri, T., Nichols, B., Quince, C., and Mahé, F. (2016). VSEARCH: A versatile open source tool for metagenomics. PeerJ 2016, e2584. doi: 10.7717/peerj.2584.

Shehzad, W., Riaz, T., Nawaz, M. A., Miquel, C., Poillot, C., Shah, S. A., et al. (2012). Carnivore diet analysis based on next-generation sequencing: application to the leopard cat (*Prionailurus bengalensis*) in Pakistan. Mol Ecol 21, 1951–1965. doi: 10.1111/j.1365-294X.2011.05424.x.

Sousa, L. L., Silva, S. M., and Xavier, R. (2019). DNA metabarcoding in diet studies: unveiling ecological aspects in aquatic and terrestrial ecosystems. Environmental DNA 1, 199–214. doi: 10.1002/edn3.27.

Srivathsan, A., Ang, A., Vogler, A. P., and Meier, R. (2016). Fecal metagenomics for the simultaneous assessment of diet, parasites, and population genetics of an understudied primate. Front Zool 13, 1–14. doi: 10.1186/s12983-016-0150-4.

Srivathsan, A., Sha, J. C. M., Vogler, A. P., and Meier, R. (2015). Comparing the effectiveness of metagenomics and metabarcoding for diet analysis of a leaf-feeding monkey (Pygathrix nemaeus). Mol Ecol Resour 15, 250–261. doi: 10.1111/1755-0998.12302.

Strum, S. C. (2005). Measuring success in primate translocation: A baboon case study. Am J Primatol 65, 117–140. doi: 10.1002/AJP.20103.

Tedersoo, L., Anslan, S., Bahram, M., Põlme, S., Riit, T., Liiv, I., et al. (2015). Shotgun metagenomes and multiple primer pair-barcode combinations of amplicons reveal biases in metabarcoding analyses of fungi. MycoKeys 10, 1–43. doi: 10.3897/mycokeys.10.4852.

Turpie, J. K., Adams, J. B., Joubert, A., Harrison, T. D., Colloty, B. M., Maree, R. C., et al. (2002). Assessment of the conservation priority status of South African estuaries for use in management and water allocation. Water SA 28.

van Wassenbergh, S., Strother, J. A., Flammang, B. E., Ferry-Graham, L. A., and Aerts, P. (2008). Extremely fast prey capture in pipefish is powered by elastic recoil. J R Soc Interface 5, 285–296. doi: 10.1098/rsif.2007.1124.

Weiss, S.-E., Emami-Khoyi, A., Kaiser, H., Cowley, P. D., James, N. C., Jansen van Vuuren, B., et al. (2022). The last two remaining populations of the Critically Endangered estuarine pipefish are inbred and not genetically distinct. Front Mar Sci 8, 1–17. doi: 10.3389/fmars.2021.756595.

Whitfield, A. K. (1995). Threatened fishes of the world: *Syngnathus watermeyeri* Smith, 1963 (Syngnathidae). Environ Biol Fishes 43, 1–2. doi: 10.1007/BF00002484.

Whitfield, A. K., Mkare, T. K., Teske, P. R., James, N. C., and Cowley, P. D. (2017). Life-histories explain the conservation status of two estuary-associated pipefishes. Biol Conserv 212, 256–264. doi: 10.1016/j.biocon.2017.06.024.

Wickham, H. (2016). ggpolt2: Elegant graphics for data analysis. 2nd ed. Springer-Verlag New York.

Wooldridge, T. H., and Callahan, R. (2000). The effects of a single freshwater release into the Kromme Estuary. 3: Estuarine zooplankton response. Water SA 26, 311.

